# NDEx IQuery: a multi-method network gene set analysis leveraging the Network Data Exchange

**DOI:** 10.1101/2022.10.24.513552

**Authors:** RT Pillich, J Chen, C Churas, D Fong, T Ideker, SN Liu, BM Gyori, K Karis, K Ono, A Pico, D Pratt

## Abstract

**Motivation:** The investigation of sets of genes using biological pathways is a common task for researchers and is supported by a wide variety of software tools. This type of analysis generates hypotheses about the biological processes active or modulated in a specific experimental context.

**Results:** The NDEx Integrated Query (IQuery) is a new tool for network and pathway-based gene set interpretation that complements or extends existing resources. It combines novel sources of pathways, integration with Cytoscape, and the ability to store and share analysis results. The IQuery web application performs multiple gene set analyses based on diverse pathways and networks stored in NDEx. These include curated pathways from WikiPathways and SIGNOR, published pathway figures from the last 27 years, machine-assembled networks using the INDRA system, and the new NCI-PID v2.0, an updated version of the popular NCI Pathway Interaction Database. IQuery’s integration with MSigDB and cBioPortal now provides pathway analysis in the context of these two resources.

**Availability and Implementation:** IQuery is available at https://www.ndexbio.org/iquery and is implemented in Javascript and Java.

**Contact:** Dexter Pratt

## Introduction

Investigating sets of proteins and genes of interest using biological pathways and networks is a common task for researchers. Networks describe relationships between genes, proteins, small molecules, and other entities and express the mechanisms associated with a biological process or the function of a cellular component. Pathway resources such as KEGG (Okuda et al. 2008), Reactome (Fabregat et al. 2015), and SIGNOR (Perfetto et al. 2016) go beyond ontology-based enrichment sources (Reimand et al. 2007; Sherman et al. 2007; Kuleshov et al. 2019) by providing information about the relationships in a curated set of genes. Other gene set analysis resources such as STRING (Szklarczyk et al. 2021), GeneMania (Warde-Farley et al. 2010), and IntAct (Del Toro et al. 2022) consult databases of molecular interactions and find relationships involving the queried genes.

This rich landscape of tools serves the biomedical research community well, but unmet needs remain. First, many sources of molecular interaction data are not aggregated and cannot be queried or have limited query interfaces. Second, where network content is aggregated, it is integrated into a single database with a common schema. While this benefits consistency, it nevertheless loses the data’s original structure and can obscure qualitative differences between sources. This normalization process also increases the effort to add and update content, slowing the pace at which databases can grow and making community contributions more complex. Third, the diversity of sources poses challenges for the reuse of query results in subsequent analyses, each source exporting in a different format or accessed via a different API. Last but not least, sharing and publishing analysis results in a computable and editable format is difficult, and most result networks are published and presented only as static images in figures. Here we present IQuery, a new tool for network and pathway-based gene set interpretation. IQuery addresses the unmet needs described above, providing functionality that complements or extends existing resources. It combines novel sources of pathways/networks, and its integration with the Network Data Exchange (NDEx) (Pratt et al. 2015; Pratt et al. 2017; Pillich et al. 2021) provides the capability to store and share analysis results.

## System and Methods

### Resources and code availability

The IQuery web user interface is implemented in Javascript using the React framework. The IQuery web application is available at https://www.ndexbio.org/iquery.

The IQuery service is implemented in Java. The source code and documentation are available on GitHub at https://github.com/ndexbio/iquery.

To use IQuery, visit https://www.ndexbio.org/iquery using any major web browser.

To report a bug, please use the contact form available at https://home.ndexbio.org/report-a-bug/. For data availability, see the Acknowledgements section.

### Cytoscape Exchange (CX) network format

CX is a JSON-based network exchange format, a flexible structure for the transmission of networks. It is designed for flexibility, modularity, and extensibility and as a message payload in common REpresentational State Transfer (REST) protocols. It is not intended as an in-memory data model for use in applications. The flexibility of CX enables straightforward strategies for lossless encoding of potentially any network. At the most basic level, this means that CX imposes very few restrictions: networks can be cyclic or acyclic, and edges are implicitly directed, but networks can use custom annotation schemes to override this. CX does not make any commitment to a single “correct” model of biology or graphic markup scheme. CX is designed to facilitate streaming, potentially reducing the memory footprint burden on applications processing large CX networks.

### Methods for ranking query results

IQuery calculates measures of alignment of a query gene set against a large collection of networks and then returns ranked lists of results. To create this ranking, IQuery implements three different metrics, each highlighting a different aspect of alignment: similarity, p-value, and overlap.

#### Similarity: Cosine similarity of the query genes and the network genes

This score characterizes the similarity between the query set and the genes in the network while considering that some genes are much more universal than others and will appear in many more networks. The cosine similarity calculation uses values derived from each gene’s term frequency-inverse document frequency (TF-IDF) in the query set and the network. Rare genes that are shared between the query set and the network will contribute more to the similarity score than common genes, resulting in a higher similarity score. When sorting by similarity, networks with high similarity are at the top of the list, and networks with low similarity are at the bottom of the list.

#### P-Value: Hypergeometric test adjusted for false discovery

This score is the probability that the query set and the network overlap, calculated using the complementary cumulative distribution function of a hypergeometric distribution, where:

- The population size (N) is equal to the number of genes in the database (i.e., the total number of unique genes found in all the pathways for any given analysis tab);
- The number of success states in the population (K) is equal to the number of genes in the network;
- The sample size (n) is equal to the size of the network; and
- The number of observed successes (k) is equal to the number of genes that are in both the query set and the network.

The p-values are adjusted to compensate for the high false discovery rate that is an effect of querying a large database of networks. This is done using the Benjamini-Hochberg method, where each p-value is multiplied by the number of networks queried and then divided by its rank relative to other p-values (where low p-values have a low rank and vice versa). When sorting by p-value, networks with a low p-value are at the top of the list, and networks with a high p-value are at the bottom of the list.

#### Overlap: the number of genes that are in both the query set and the network

When sorting by overlap, networks with a high number of overlapping genes are at the top of the list, and networks with a low number of overlapping genes are at the bottom of the list. Sorting is stable, so sorting by p-value and then by overlap will result in a list where networks are sorted by overlap, and networks that are tied for overlap are sorted by p-value.

## Relationship to other resources

IQuery is part of the Network Data Exchange (NDEx) infrastructure (https://www.ndexbio.org/). It draws on curated pathway content from WikiPathways (https://www.wikipathways.org/) and the SIGNOR database (https://signor.uniroma2.it/), as well as machine-assembled networks from the INDRA system (https://www.indra.bio/) and published pathway diagrams from the Pathway Figures OCR (https://pfocr.wikipathways.org/) open science project. IQuery is also integrated with the popular Molecular Signatures Database (MSigDB) (https://gsea-msigdb.org) and cBioPortal for Cancer Genomics (https://www.cbioportal.org/).

## Results

### User interface

The IQuery interface has several elements, as shown by the colored outlines in Figure 1, centered on the graphical rendering module identified by the orange outline.

**Figure 1.**
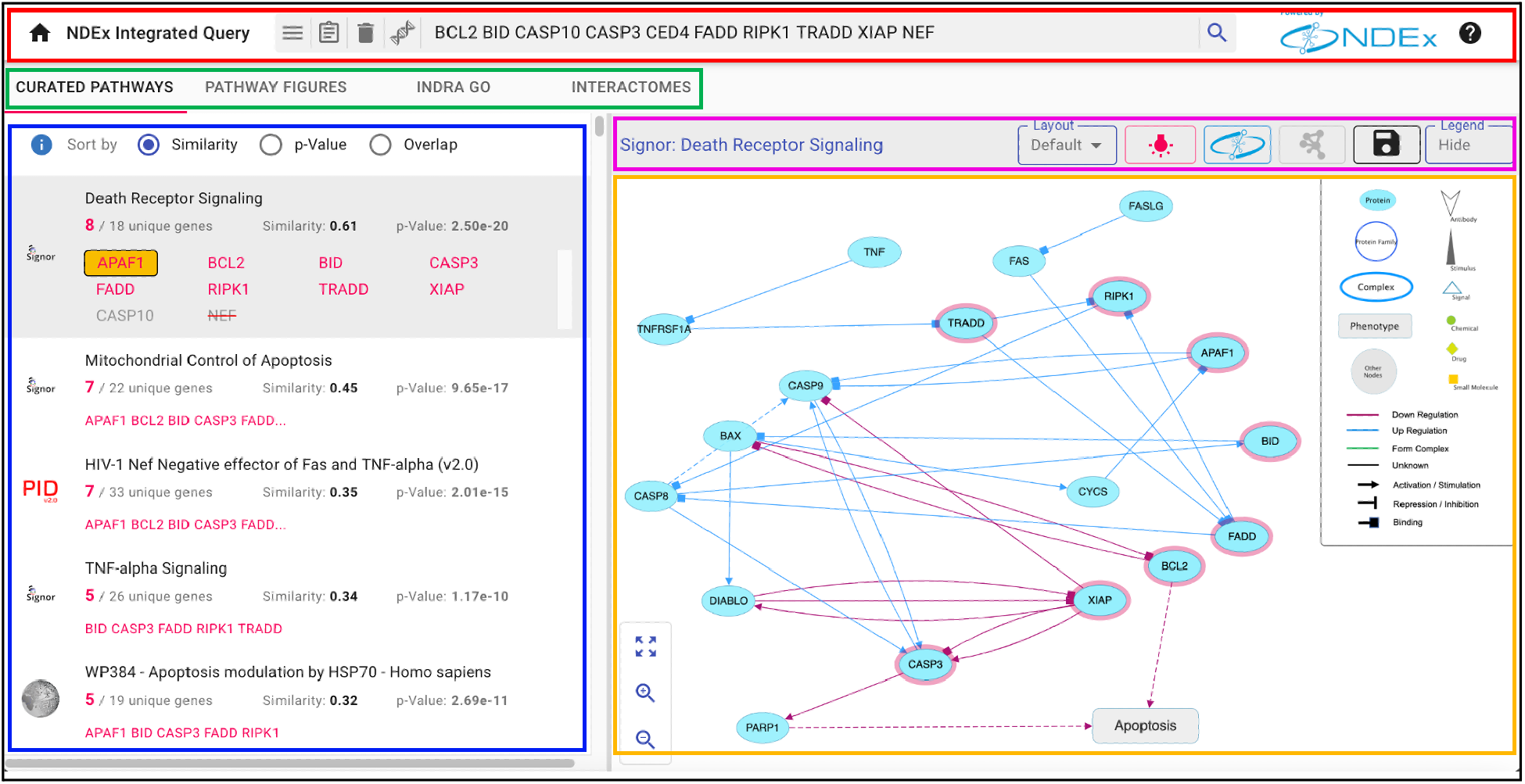
The Curated Pathways tab results for an apoptosis-related set of 10 genes (BCL2 BID CASP10 CASP3 CED4 FADD RIPK1 TRADD XIAP NEF). The top result displayed is the Death Receptor Signaling pathway from the SIGNOR database. Additional results include NCI-PID v2.0 and WikiPathways networks. For each result, matched query genes are shown in red, normalized genes boxed in orange and invalid genes with a strikethrough. Red outline: Input element. Green outline: Analysis element. Blue outline: Results element. Purple outline: Toolbar element. Orange outline: Graphical Rendering element.

#### Input Element

The input element (red outline) provides a search box to input query genes, augmented with buttons to clear the query, view it in full or select one of the available example gene sets. On the right-hand side, the black “question mark” icon lets users access a short tutorial on the interface and its modules.

#### Analysis Element

The analysis tabs (green outline) select between the four currently supported analyses described in the analysis section below.

#### Results Element

The results panel (blue outline) displays the query results of each analysis along with details about the matched query genes.

At the top of the panel, the “Sort by” radio buttons let users choose among three different ordering methods:

1. <u>Similarity:</u> the cosine similarity of the query genes and the network genes.
2. <u>P-value</u>: the hypergeometric test adjusted for false discovery.
3. <u>Overlap</u>: the number of genes that are in both the query set and the network.

Additional details about the three methods are available by clicking the blue info icon next to the “Sort by” controls.

Each query result displays the number of matched genes and the number of genes in the network. When selected, the result expands to show all the query genes with matched genes highlighted in red. If the original gene set includes invalid or unofficial gene names, IQuery will normalize those names and provide details about the normalization performed.

#### Toolbar Element

The toolbar (purple outline) shows the name of the selected query result and provides common controls to perform several actions. The network’s name can be clicked in the toolbar to display a window showing its description, reference, and other attributes. This network information window also includes tabs to browse the network’s nodes and edges in tabular form. The same tabs are also displayed when individual nodes and edges are selected in the graphical rendering to explore their annotations. Annotations are specific to the various types of networks and can include information such as publications supporting an interaction. Other toolbar controls let users change the network’s layout, toggle the highlighting of query genes, and display the network’s legend. Finally, two additional dedicated toolbar buttons integrate IQuery with Cytoscape and NDEx.

#### Integration with Cytoscape

IQuery is tightly integrated with Cytoscape (Shannon et al. 2003), the widely used network visualization and analysis application, to enable immediate use of query results in further analyses. IQuery result networks can be seamlessly downloaded to Cytoscape to be edited, used to generate publication-quality images, merged with other networks, annotated with user datasets, or used with any of hundreds of community-developed apps (Lotia et al. 2013).

#### Integration with NDEx to store and share results

NDEx, the Network Data Exchange (Pratt et al. 2015; Pratt et al. 2017; Pillich et al. 2021), is the storage and sharing cloud component used by Cytoscape and its umbrella of tools and services. In the NDEx web interface, users can manage their networks in everyday work, make them publicly available, and link to them from publications. IQuery enables users to save query results to their NDEx account, complete with node annotations indicating the query genes.

### Analysis

The IQuery web application performs four separate gene set analyses based on a diverse range of pathways/networks from NDEx and presents the results in four dedicated tabs: Curated Pathways, Pathway Figures, INDRA GO, and Interactomes.

#### Curated Pathways

In this tab, IQuery aggregates curated pathway content from multiple sources. WikiPathways (Martens et al. 2021) is an open, collaborative, community-driven platform dedicated to the curation of biological pathways. The pathway models were contributed, annotated, and updated by over 700 individuals over the past two decades. Community curators review every edit prior to its inclusion in the official monthly release. IQuery performs custom gene set analysis on a suite of 675 curated human pathways and 11 cancer hallmark networks, each of which is an aggregation of between 3 and 18 pathway models categorized by the Clinical Proteomic Tumor Analysis Consortium (CPTAC) (https://cptac.wikipathways.org).

SIGNOR (Perfetto et al. 2016) is a repository of manually annotated experimental evidence about causal interactions between proteins and other entities of biological relevance. SIGNOR includes signaling, metabolic, disease, and cancer pathways as well as COVID-19 network hallmarks, providing a total of 90 models to IQuery.

NCI-PID v2.0 is an updated version of the Pathway Interaction Database (PID) (Schaefer et al. 2009). The PID was created in a collaboration between the US National Cancer Institute and Nature Publishing Group in 2009 and was last updated in 2012. As the maintainers of this popular resource, we are now augmenting the original pathways with machine-generated relationships on an ongoing basis, capturing the latest, up-to-date information. These relationships are assembled using the INDRA system (Gyori et al. 2017) and are annotated with links to detailed summaries of supporting literature evidence, including the specific supporting text. INDRA is described in detail in the INDRA GO analysis section below.

In the Curated Pathways tab, all protein node names are normalized to official gene symbols but generally preserve the semantics and curation approaches intended by the authors. For example, the WikiPathways models retain their diagram format and heterogeneous visual styling features as expected from a community-curated resource, while the SIGNOR pathways keep their causal evidence edge annotations.

#### Pathway Figures

Drawing on the Pathway Figure OCR open science project (Hanspers et al. 2020), IQuery includes a collection of over 30,000 gene sets defined by published pathway figures. Using machine learning, they first identified 80,000 pathway figures in PMC-indexed articles between 1995 and 2021, and then used optical character recognition (OCR) and named entity recognition (NER) to identify genes, proteins, chemicals, and diseases mentioned in each figure. Altogether, 1.5 million genes (14,253 unique genes) were identified in published network and pathway figures, representing more breadth and depth of content than a typical pathway database. The resulting machine-readable gene sets offer a novel means of analyzing published pathway models. The IQuery collection is limited to the 32,263 gene sets containing six or more genes to optimize for query similarity, enrichment, and overlap. Each set is represented as an edgeless network and is presented along with the original figure and parent article citation. In IQuery, this corpus enables users to find pathways specific to individual articles, providing detailed biological and experimental context for interpreting their query results (Figure 2). Moreover, genes and processes that are not well represented in public curated collections may be found in published pathway figures reflecting recent findings or less-studied topics.

**Figure 2.**
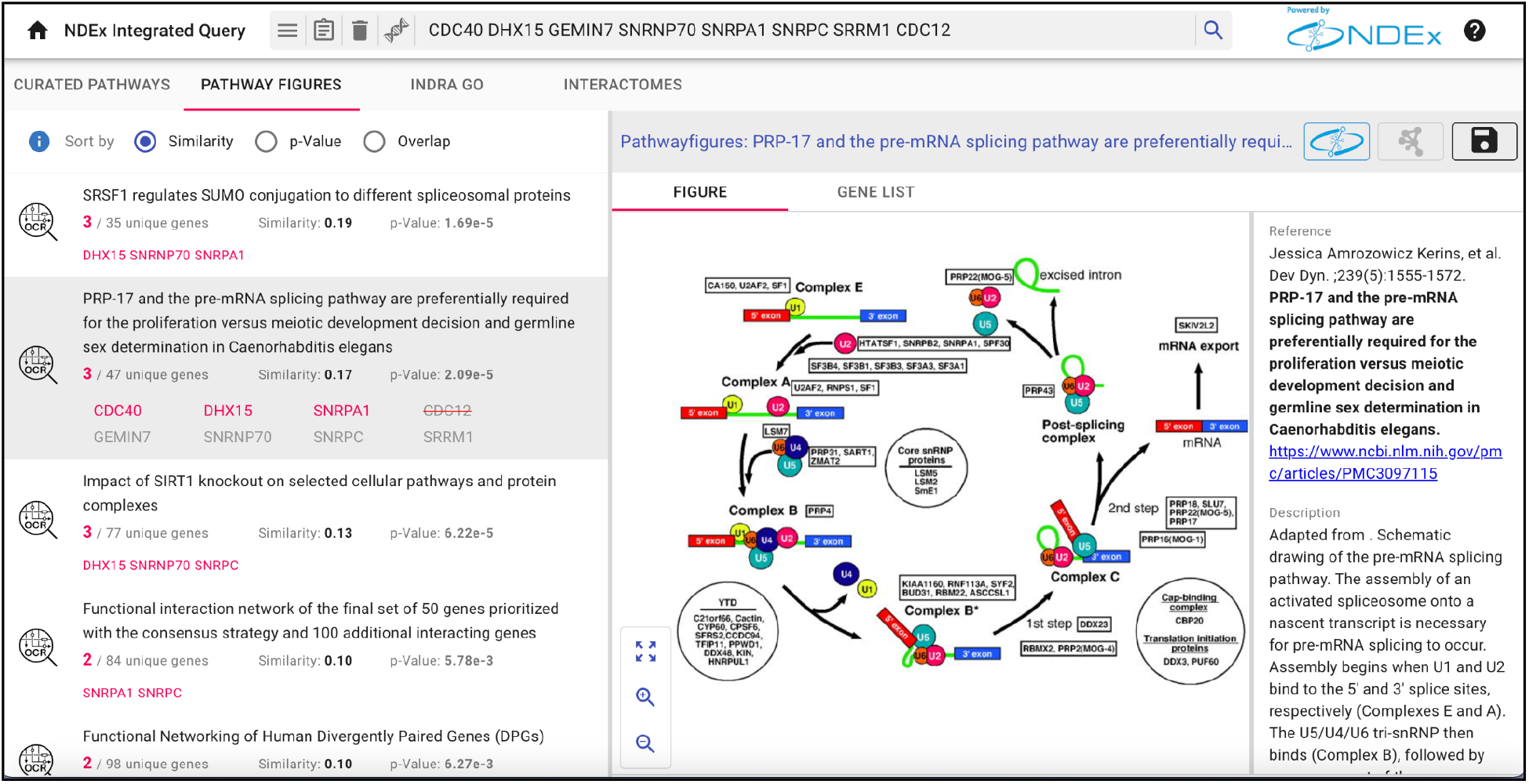
(A) An mRNA processing-related gene set (CDC12 CDC40 DHX15 GEMIN7 SNRNP70 SNRPA1 SNRPC SRRM1) finds pathway figures from papers specifically focused on the spliceosome.

#### INDRA GO

We used the INDRA system to assemble the output of multiple automated literature mining systems (Valenzuela-Escárcega et al. 2018) (which were run via INDRA on PubMed abstracts and PubMedCentral full-text articles to extract regulation and interaction among biological entities) with the content of structured pathway knowledge bases including Pathway Commons (Rodchenkov et al. 2019). We then used this assembled pathway knowledge base to create networks of interactions and regulations among the set of genes annotated with a given GO term. The INDRA GO tab displays the results of queries against a set of 6,295 networks created for small (< 200 genes) GO Biological Process terms. Edges between genes in these networks summarize the high-confidence relationships found by INDRA. As in the case of NCI-PID v2.0 described above, the INDRA-derived relationships are up-to-date and have links to detailed summaries of supporting literature evidence, including the specific supporting text (Figure 3).

**Figure 3.**
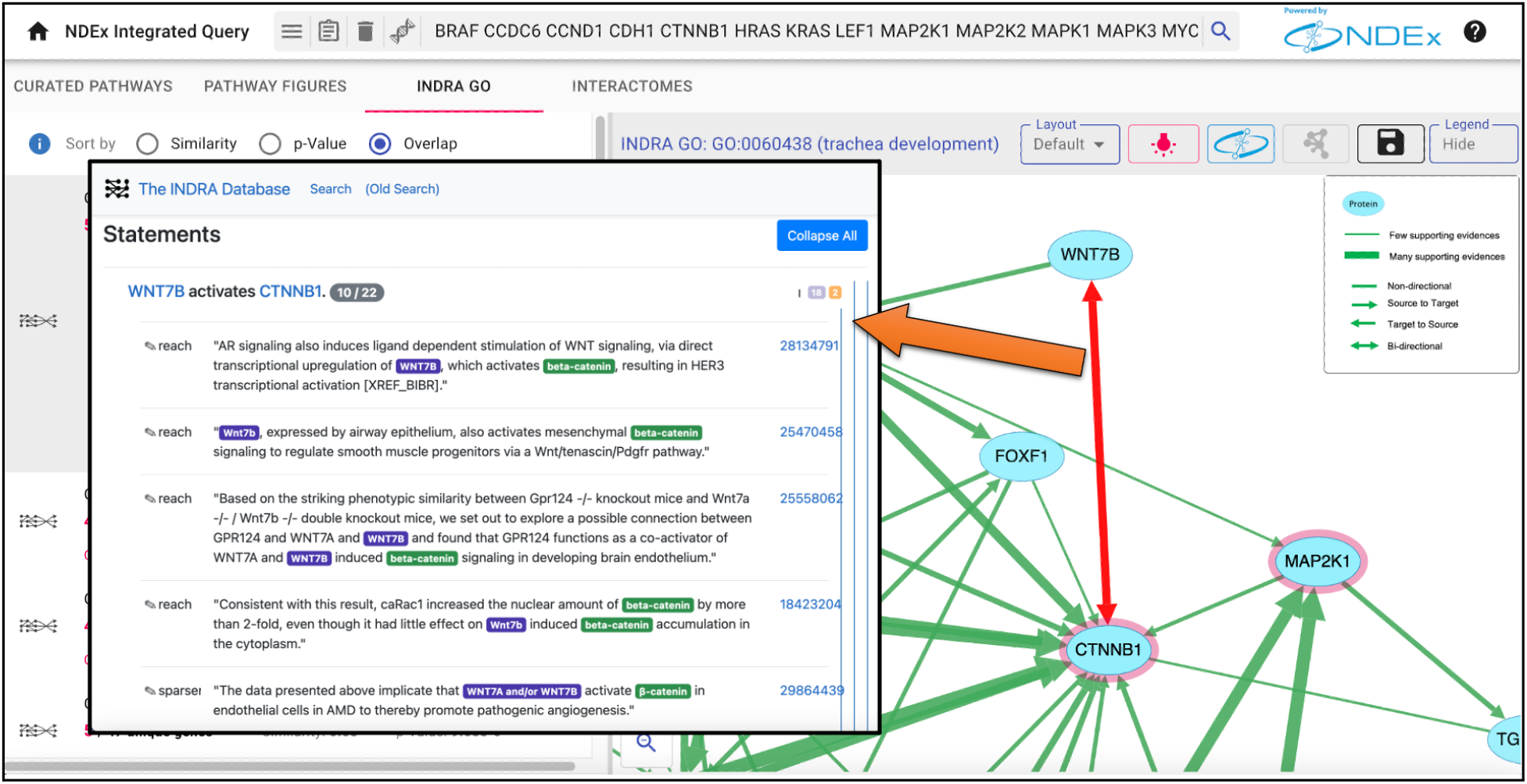
The top-ranking INDRA GO network for an input gene set is the GO term for trachea development. The network edges represent interactions or regulations assembled by INDRA among genes annotated with this GO term, including WNT7B and CTNNB1. The evidence supporting the relationship between WNT7B and CTNNB1 can be examined in detail by clicking on the edge, which links to the INDRA Database, where specific evidence sentences and links to source publications are provided.

#### Interactomes

In the Interactomes tab, users can run on-demand queries on selected large interaction networks stored in NDEx (Pratt et al. 2015; Pratt et al. 2017; Pillich et al. 2021). As of version 1.4, IQuery uses 20 interactomes derived from public databases or published by researchers. One of the interactomes is BioPlex 3 (Huttlin et al. 2021), the latest version of the high-quality network assembled from protein-protein interactions identified in both HEK293T and HCT116 cells by affinity purification and mass spectrometry. Another example is the human protein interaction network from the BioGRID database (Oughtred et al. 2021). IQuery and NDEx (Pratt et al. 2015; Pratt et al. 2017; Pillich et al. 2021) provide convenient access to these datasets, complementing the single-gene query capability of their own user interfaces. In addition, many interaction networks used in IQuery are published by researchers. In these cases, IQuery allows easy access and use of data that might otherwise be relegated to supplemental materials, not accessible by user queries. The type of query and the specific target interaction network to analyze are user-defined, and the analysis is only run if the user decides to do so. This makes IQuery fast and efficient while preserving the flexibility to adapt to individual use cases. The query results are opened in the NDEx web interface in a new browser tab; here, users can refine or modify their query, save their result, or open it in Cytoscape (Shannon et al. 2003) for further analysis.

### Integration

IQuery has a modular, service-oriented structure in which a primary service aggregates other services. The IQuery service accepts query strings and returns (1) lists of networks as query results and (2) networks in CX format (see System and Methods). The integration of IQuery into a web application is simple. It can be embedded in the application via the “IFRAME” HTMLtag and passed gene lists via an HTTP “GET” transaction (Figure 4A). Alternatively, the application can open IQuery in a new tab. This method was developed in a collaboration with The Molecular Signatures Database (MSigDB) (Liberzon et al. 2015). MSigDB is a widely used and comprehensive database of over 10,000 gene sets for performing gene set enrichment analysis. MSigDB users can now send any gene set to IQuery for further analysis. A collaboration between The cBioPortal for Cancer Genomics (Gao et al. 2013) and the NDEx project is an excellent example of advanced IQuery integration. cBioPortal provides a web resource for exploring, visualizing, and analyzing multidimensional cancer genomics data. The cBioPortal query interface combined with customized data storage enables researchers to interactively explore genetic alterations across samples, genes, and pathways. A cancer-focused version of IQuery was embedded in the cBioPortal interface, enhanced to enable users to view their genes of interest annotated with genetic alteration frequency data from selected datasets (Figure 4B).

**Figure 4.**
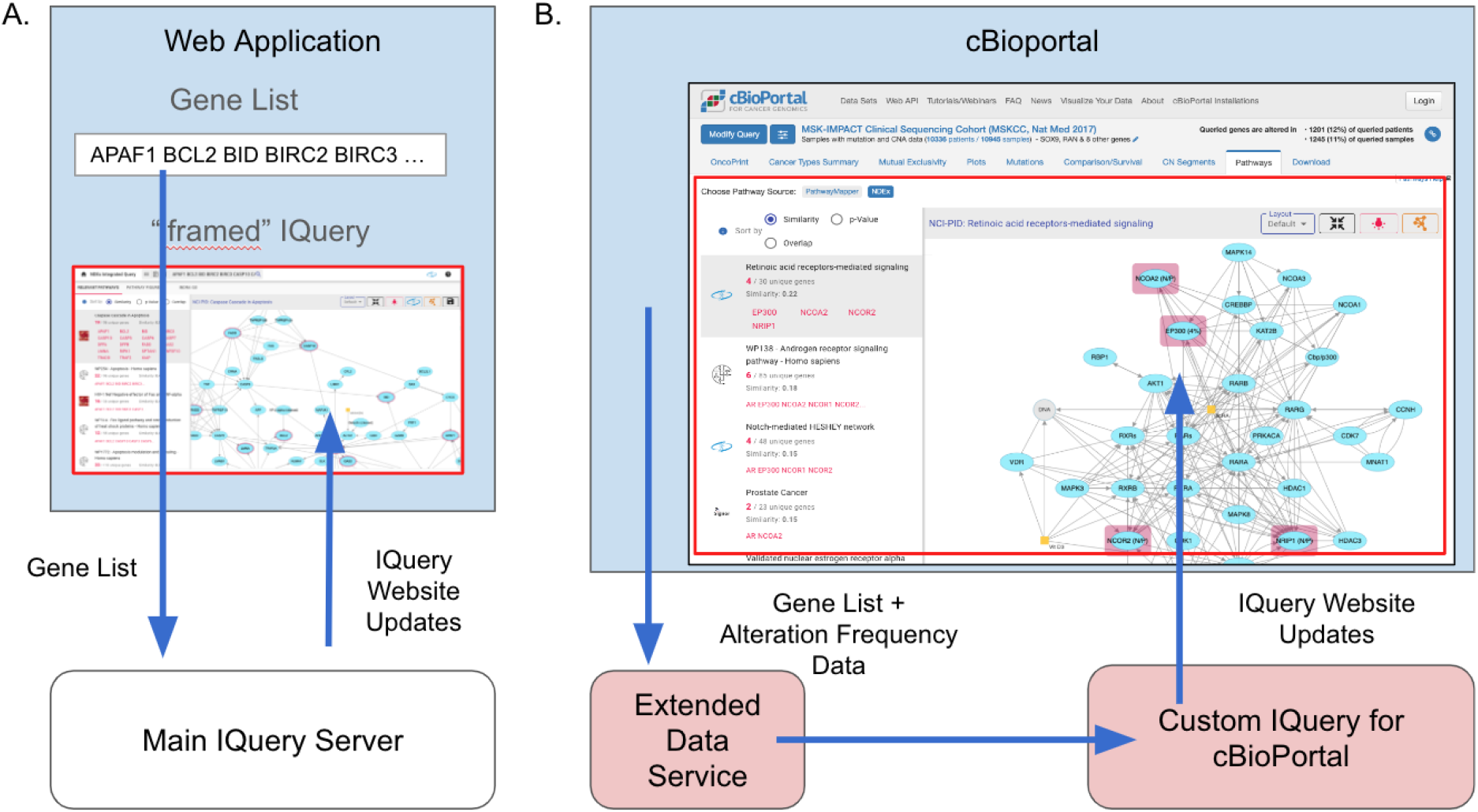
Web application integration of IQuery. (A) Basic integration of IQuery. (B.) Advanced Query integration with cBioPortal.

The alteration frequency data is passed to the IQuery server via an extended data service endpoint.

## Discussion

### Interpreting IQuery results

Given the diversity of the pathway networks on which IQuery operates, the values of the scores for a given pathway or its rank in the results should be seen as general advice, a means of directing the user’s attention to potentially relevant mechanisms. Pathways relevant to a given mechanism from different sources reflect alternative choices of genes and may significantly differ in size. This can provide complementary information about the relationships of the genes and proteins within the pathways but can also affect the ranking of those pathways. For example, a gene set related to Apoptosis finds pathways of various sizes from SIGNOR (Perfetto et al. 2016), NCI-PID 2.0, and WikiPathways (Martens et al. 2021). A larger pathway may be more informative than a smaller pathway with a better p-value score if it contains more overlapping genes or if the proteins are clustered in a small subnetwork.

### Machine-driven pathway augmentation

With NCI-PID v2.0, we took the first step toward a sophisticated, machine-driven approach to refresh an outdated pathway resource, augmenting the original PID (Schaefer et al. 2009) pathways with the addition of up-to-date relationships. However, the proteins, chemicals, and complexes selected for inclusion in each pathway must also be updated to reflect the new research findings generated in the last ten years. In the next step, we will generate a new pipeline that combines both the INDRA (Gyori et al. 2017) and Pathway Figure OCR (Hanspers et al. 2020) technologies to expand the pathways with additional entities relevant to the mechanisms and current members. In this way, the NCI-PID will continue to grow and be maintained as a unique, dynamic reference resource.

### Embedded IQuery and community-sourced data

The next steps with IQuery will focus on (1) expansion via community-sourced pathway and interaction data, (2) new collaborations in which IQuery is embedded in applications, and (3) enabling organizations to operate IQuery servers focused on their own data. IQuery provides collaborators with a simple way to distribute their experimentally derived content and expose it to the community for immediate validation and reuse. Projects that output large datasets, such as BioPlex (Huttlin et al. 2021), HIPPIE (Alanis-Lobato et al. 2017), or ProteomeHD (Kustatscher et al. 2019), can rely on the mature NDEx (Pratt et al. 2015; Pratt et al. 2017; Pillich et al. 2021) infrastructure to publish their data, rather than developing their own web interface or database. The networks used in the analyses can be customized when embedding IQuery in an application, such as the networks produced by a specific lab. The ability of organizations to deploy private instances of IQuery and NDEx enables the use of IQuery with protected or proprietary data behind firewalls.

To conclude, IQuery is a flexible framework for pathway and interaction network-based analysis of gene sets; it encourages collaborations within the research community while providing a channel for researchers to make their network models immediately available and usable by other scientists.

## Data availability

All data sources used by IQuery are publicly available and indexed for search in the Network Data Exchange (NDEx) at https://www.ndexbio.org. In addition, the two novel data sources described in the manuscript, NCI-PID v2.0 and INDRA GO, are also accessible via DOI at https://doi.org/10.18119/N93W46 and https://doi.org/10.18119/N9060V respectively.

## References

Alanis-Lobato, G. et al. (2017). HIPPIE v2.0: enhancing meaningfulness and reliability of protein-protein interaction networks. Nucleic Acids Res. 45, D408–414

Del Toro, N. et al. (2022). The IntAct database: efficient access to fine-grained molecular interaction data. Nucleic Acids Res. 50, D648–653

Fabregat, A. et al. (2015). The Reactome pathway Knowledgebase. Nucleic Acids Res. 44, D481–487

Gao, J. et al. (2013). Integrative analysis of complex cancer genomics and clinical profiles using the cBioPortal. Sci. Signal. 6 (269), pl1

Gyori, B. M. et al. (2017). From word models to executable models of signaling networks using automated assembly. Mol Syst Biol. 13(11), 954.

Hanspers, K. et al. ((2020). Pathway information extracted from 25 years of pathway figures. Genome Biol 21, 273

Huttlin, E. L. et al. (2021). Dual proteome-scale networks reveal cell-specific remodeling of the human interactome. Cell 184, 3022–3040.

Kuleshov, M. V. et al. (2019). modEnrichr: a suite of gene set enrichment analysis tools for model organisms. Nucleic Acids Res. 47, W183–190

Kustatscher, G. et al. (2019). Co-regulation map of the human proteome enables identification of protein functions. Nat Biotechnol. 11, 1361–1371.

Liberzon, A. et al. (2015). The Molecular Signatures Database (MSigDB) hallmark gene set collection. Cell Syst. 1, 417–425

Lotia, S. et al. (2013). Cytoscape app store. Bioinformatics 29, 1350–1351

Martens, M. et al. (2021). WikiPathways: connecting communities. Nucleic Acids Res. 49, D613–621

Okuda, S. et al. (2008). KEGG Atlas mapping for global analysis of metabolic pathways. Nucleic Acids Res. 36, W423–426

Oughtred, R. et al. (2021). The BioGRID database: A comprehensive biomedical resource of curated protein, genetic, and chemical interactions. Protein Sci. (1), 187–200

Perfetto, L. et al. (2016). SIGNOR: a database of causal relationships between biological entities. Nucleic Acids Res. 44, D548–554

Pillich, R. T. et al. (2021). NDEx: Accessing Network Models and Streamlining Network Biology Workflows. Curr Protoc. 1(9), e258

Pratt, D. et al. (2015). NDEx, the Network Data Exchange. Cell Syst. 1(4), 302–305

Pratt, D. et al. (2017). NDEx 2.0: A Clearinghouse for Research on Cancer Pathways. Cancer Res. 77(21), e58–e61

Reimand, J. et al. (2007). g:Profiler - a web-based toolset for functional profiling of gene lists from large-scale experiments. Nucleic Acids Res. 35, W193–200

Rodchenkov, I. et al. (2019). Pathway Commons 2019 Update: integration, analysis and exploration of pathway data. Nucleic Acids Res. 48, D489–497.

Schaefer, C.F. et al. (2009). PID: the Pathway Interaction Database. Nucleic Acids Res. 37, D674–679

Shannon, P. et al. (2003). Cytoscape: a software environment for integrated models of biomolecular interaction networks. Genome Res. 13, 2498–2504

Sherman, B. T et al. (2007). DAVID Knowledgebase: a gene-centered database integrating heterogeneous gene annotation resources to facilitate high-throughput gene functional analysis. BMC Bioinformatics 8, 426

Szklarczyk, D. et al. (2021). The STRING database in 2021: customizable protein-protein networks, and functional characterization of user-uploaded gene/measurement sets. Nucleic Acids Res. 49, D605–612

Valenzuela-Escárcega, M.A. et al. (2018). Large-scale automated machine reading discovers new cancer-driving mechanisms. Database (Oxford). 2018:bay098

Warde-Farley, D. et al. (2010). The GeneMANIA prediction server: biological network integration for gene prioritization and predicting gene function. Nucleic Acids Res. 38, W214–220

